# OmniCell: Unified Foundation Modelling of Single-Cell and Spatial Transcriptomics for Cellular and Molecular Insights

**DOI:** 10.64898/2025.12.29.696804

**Authors:** Jiangshuan Pang, Ping Qiu, Youzhe He, Yiting Deng, Wangyang Tang, Zhi Huang, Jing Yan, Baolong Li, Adi Lin, Lei Cao, Fei Teng, Shuangsang Fang, Shengkang Li, Ziqing Deng, Yong Zhang, Yuxiang Li, Shaoshuai Li, Xun Xu

**Author notes:** Corresponding authors: Email addresses (Yong Zhang), (Yuxiang Li), (Shaoshuai Li), (Xun Xu). These authors contributed equally to this work (Jiangshuan Pang, Ping Qiu, Youzhe He, Yiting Deng, Wangyang Tang).

## Abstract

A cell’s transcriptional programme is not fully defined by gene expression alone, but by the tissue context in which that programme is enacted. Single-cell RNA sequencing resolves molecular identity after dissociation, whereas spatial transcriptomics preserves tissue architecture but remains constrained by assay-specific sparsity and gene coverage. Here we present OmniCell, a tissue-contextual transcriptomic foundation model pretrained on 67 million dissociated and spatially resolved profiles. By integrating gene identity, expression magnitude and tissue context, OmniCell links transcriptional programmes to the cellular neighbourhoods and anatomical contexts in which they operate. OmniCell organised transcriptomes across molecular, cellular and tissue scales. It recovered cell-type-specific programmes and tissue-aligned gene modules, preserved robust cell-state structure across batches, species and rare populations, and improved the reconstruction of spatial cell identity, anatomical domains and cell-type composition. In human liver cancer Stereo-seq data, OmniCell resolved a tumour-margin transition zone characterised by immune infiltration, acute-phase inflammation, coagulation/complement activity and metallothionein-linked metal-ion detoxification. Contextual gene-embedding similarity analysis showed that gene relationships differed across tumour core, transition-zone and paratumour/adjacent non-malignant niches, indicating that OmniCell captures tissue-dependent gene function rather than expression similarity alone. In mouse brain development and macaque cortex, spatial virtual perturbations mapped regulatory genes onto stage- and region-specific anatomical programmes. Together, these results establish tissue context as a primary axis of transcriptomic representation and provide a framework for studying how cellular programmes acquire context-dependent biological meaning in intact tissues.

## 1. Introduction

A cell’s molecular profile is only part of its biological identity. The same transcriptional programme can have different biological meaning at an invasive tumour border, within an immune-rich niche or in normal parenchyma. In each setting, cellular state is shaped not only by lineage and intrinsic regulation, but also by the surrounding tissue environment^1^. Single-cell RNA sequencing (scRNA-seq) has transformed cellular cataloguing across organs, diseases and species^2–8^, but dissociation removes the native coordinates, cell-cell contacts and microenvironmental cues that often determine how a programme should be interpreted^2–4,9^. Spatial transcriptomic technologies restore this address by measuring RNA in intact tissue, from imaging-based assays such as MERFISH, seqFISH and STARmap to in situ sequencing and spatial barcoding platforms such as Visium and Slide-seq^10–15^. Computational spatial methods now further support cell localisation, tissue-module discovery and the modelling of spatial gene-expression variation in high-resolution data^16–18^. These developments have made tissue architecture increasingly accessible to transcriptomic analysis, but they do not remove the need for representations that can connect molecular depth with anatomical context. Dissociated single-cell atlases capture rich cellular states but lose native tissue organisation, whereas spatial assays preserve anatomical structure with platform-dependent sparsity, resolution and gene coverage^2–4^. This creates a central challenge for transcriptomic representation learning: how to model cellular programmes together with the tissue environments in which they operate.

Transcriptomic foundation models offer a way to learn reusable structure from heterogeneous single-cell data. Models such as scGPT, scFoundation and Geneformer have shown that gene-expression profiles can be represented as structured molecular sequences, producing embeddings that support annotation, perturbation analysis and other downstream tasks^19–22^. This shift has moved the field from task-specific pipelines toward general cellular representations. However, most existing transcriptomic foundation models are trained primarily on dissociated cell atlases. They learn the grammar of gene expression and cell identity, but they do not directly learn the tissue syntax through which neighbourhood, anatomy and microenvironment constrain cell behaviour. For brain tissue, tumours and immune-infiltrated organs, this missing context is not auxiliary metadata; it is part of the biological signal being modelled.

Recent spatial foundation models have begun to address this gap, but they incorporate space in different and still partial ways. scGPT-spatial extends scGPT by continual pretraining on a spatial transcriptomic corpus and adds protocol-aware mixture-of-experts decoders, spatially aware sampling and neighbourhood-based reconstruction objectives^23^. These choices improve integration, deconvolution and imputation^23^, but spatial information is introduced mainly through adaptation objectives around an existing expression-language model. Nicheformer pretrains on dissociated and spatially resolved cells and evaluates spatial labels, neighbourhood composition and density prediction^24^, but its authors note that performance depends on the abundance and diversity of relevant cell and tissue types, and that explicit physical location is not incorporated during pretraining. CellPLM uses spatial transcriptomic data with two-dimensional positional encodings to learn cell-cell relations^25^, but it models cells rather than genes as tokens and evaluates spatial representation largely through imputation. These studies establish that spatial information is useful, while leaving open whether gene identity, continuous expression and observed tissue coordinates can be coupled natively during pretraining.

Here we introduce OmniCell, a tissue-contextual foundation model for single-cell and spatial transcriptomics. OmniCell was pretrained on approximately 67 million profiles, including dissociated scRNA-seq data across 45 human tissues and Stereo-seq spatial transcriptomic data across 26 tissue types. It represents each profile as a sequence of gene tokens, with each token carrying gene identity, expression magnitude and spatial position when that position is observed. For spatial transcriptomic data, OmniCell serialises a focal cell or spot together with local neighbours, allowing self-attention to operate over intracellular gene relationships and intercellular tissue context in a single computation^26^. Dissociated profiles pass through the same context-aware architecture with placeholder coordinates, sharing molecular information across modalities.

OmniCell was designed to meet three requirements for tissue-contextual transcriptomic representation. First, expression magnitude should preserve within-profile ordering while remaining robust to assay-specific scale, sequencing depth and outliers. OmniCell therefore uses soft-rank regression rather than discrete expression-bin pre-diction. Second, the same expression value should not be interpreted identically for every gene. To address this, OmniCell embeds expression magnitude through a gene-aware mixture-of-experts module that combines shared abundance trends with gene-conditioned specialisation. Third, gene activity should be interpreted in relation to the cell state and tissue neighbourhood in which it is observed. OmniCell places measured tissue coordinates directly inside self-attention through two-dimensional rotary positional encoding, and couples gene-level features with cell-level context through a symmetric bilinear output module. Together, these components allow gene identity, expression magnitude and tissue position to be learned as a single biological representation, rather than as separate molecular and spatial downstream corrections.

We assessed OmniCell along an evidence ladder that moves from representation fidelity to biological interpretation. At the cellular scale, OmniCell improved clustering and annotation across human datasets, transferred to mouse brain and muscle after homologous gene mapping, and retained cell-state structure under batch variation and simulated dropout. Its expert-routing patterns also followed lineage-associated cellular programmes. At the gene scale, contextual gene embeddings recovered canonical cell-type markers and organised genes into functional modules in both scRNA-seq and spatial transcriptomic data, including neuronal, glial and immune programmes in human Alzheimer’s disease datasets. At the tissue scale, OmniCell preserved cell identity in native spatial maps and improved spatial de-convolution and anatomical domain reconstruction^27–29^. We then tested whether this representation could expose tissue-contextual biology beyond benchmark performance. In human liver cancer Stereo-seq data, OmniCell resolved a tumourmargin transition zone marked by immune infiltration, acute-phase inflammation, coagulation/complement activity and metallothionein-linked metal-ion detoxification^1,30–31^. Contextual gene-embedding similarity analysis showed that gene relationships differed across tumour core, transition-zone and paratumour/adjacent non-malignant niches, supporting tissue-dependent gene function rather than expression similarity alone. Finally, spatial virtual perturbations of selected regulators mapped onto developmental and cortical anatomical programmes in mouse brain and macaque cortex^32–33^. Together, these analyses test whether tissue-contextual pretraining can move transcriptomic foundation modelling beyond benchmark improvement toward context-dependent gene programmes, disease-associated tissue niches and spatially organised perturbation hypotheses.

## 2. Results

### 2.1. OmniCell couples molecular identity with tissue context in a unified transcriptomic model

We first asked how a transcriptomic foundation model could represent the two biological axes that are often separated by current assays: the molecular programme of a cell and the tissue environment in which that programme is enacted. OmniCell was built to learn from both axes during pretraining rather than to add spatial information as a downstream correction. Its pretraining corpus contained approximately 67 million profiles, including dissociated scRNA-seq data and human Stereo-seq spatial transcriptomic data represented at Bin20 and Cellbin resolutions (Fig. 1a). All spatial transcriptomic datasets used for pretraining were human tissue datasets. This design exposed the model to molecularly resolved cell states and human spatial tissue contexts, providing the basis for a shared representation across single-cell and spatial assays.

**Figure 1:**
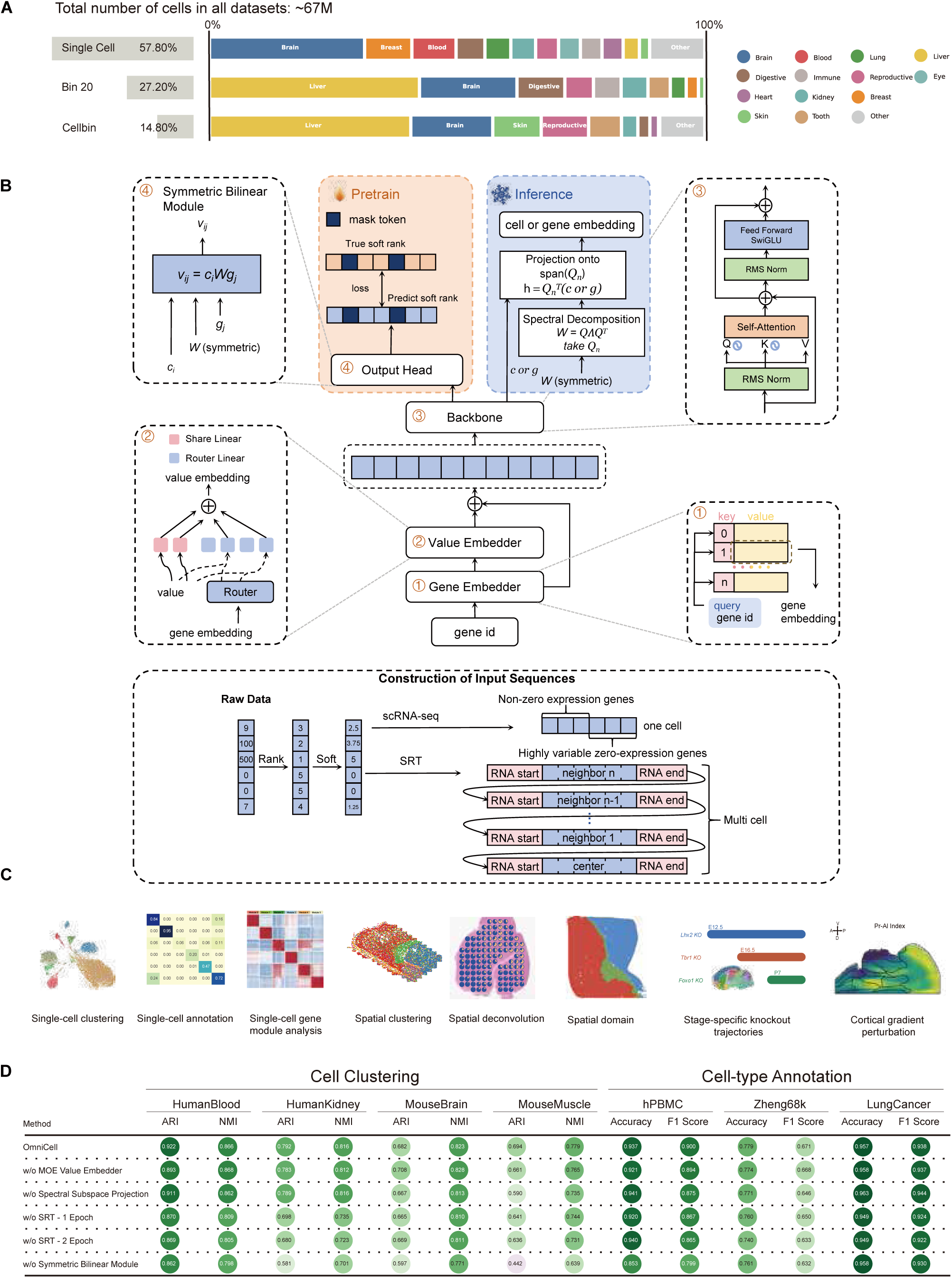
Overview of OmniCell pretraining, architecture and evaluation. (**A**) Composition of the pretraining corpus. OmniCell was pretrained on approximately 67 million profiles, including single-cell RNA-seq data (57.8%) and Stereo-seq spatial transcriptomic data represented at Bin20 (27.4%) and Cellbin (14.8%) resolutions across diverse tissue categories. (**B**) Model architecture. OmniCell encodes gene identity and expression value in a unified sequence format. Single-cell inputs prioritise non-zero expressed genes and highly variable zero-expression genes, whereas spatial inputs include a centre cell or spot and neighbouring spatial profiles. Soft-rank transformation normalises expression values, a gene-aware MoE value embedder encodes expression in gene-specific context, a Transformer backbone learns contextual representations, and a symmetric bilinear output module links cell and gene embeddings during pretraining and inference. (**C**) Downstream biological analyses enabled by the learned representation, including single-cell clustering, cell-type annotation, gene-module analysis, spatial clustering, spatial deconvolution, spatial domain identification, stage-specific knockout trajectory analysis and cortical-gradient perturbation. (**D**) Ablation analysis across cell clustering and cell-type annotation benchmarks, showing the contribution of the MoE value embedder, spectral subspace projection, spatial transcriptomic pretraining and symmetric bilinear output module.

OmniCell converts both modalities into gene-token sequences, but preserves the biological distinction between dissociated and spatial measurements (Fig. 1b). For scRNA-seq, a cell is represented by expressed genes together with highly variable zero-expression genes, allowing informative absence as well as abundance to contribute to the state. For spatial transcriptomic data, each sequence contains a focal cell or spot and its nine nearest spatial neighbours. This formulation allows self-attention to model intracellular gene relationships and local intercellular tissue context within one structured input. Continuous expression values are transformed into within-profile soft ranks, which reduce dependence on assay-specific count scale while preserving relative expression order. Gene identities and soft-rank values are then embedded through a gene-aware mixture-of-experts value embedder, enabling the same numerical expression value to be interpreted in a gene-dependent manner.

The model objective links these inputs to a tissue-contextual representation. A Transformer backbone with context-aware attention produces contextual gene and cell representations, and a symmetric bilinear output module predicts masked soft-rank values from the interaction between a cell context and a gene representation (Fig. 1b). This objective forces the model to learn whether a gene should be expressed in a particular cellular and spatial context, rather than learning a cell embedding or a gene embedding in isolation. At inference, spectral subspace projection of the learned bilinear matrix provides stable cell and gene embeddings for downstream analysis.

The resulting representation was designed to support biological questions at multiple scales (Fig. 1c). At the cell level, it can be used for clustering and annotation, and at the single-cell-to-spatial interface it can support deconvolution^27–28^. At the gene level, it supports marker discovery and gene-module analysis. At the tissue level, it enables spatial clustering, domain reconstruction and perturbation analysis along anatomical gradients^29,32–33^. We therefore treated Figure 1 not only as a model schematic, but as the entry point for the evidence ladder tested in the rest of the paper: cell-state transfer and rare-state recovery, gene-programme interpretation, spatial domain biology and spatial perturbation prediction.

Ablation experiments supported the contribution of the main architectural components (Fig. 1d). Across clustering and cell-type annotation benchmarks, the full OmniCell model generally achieved the strongest or near-strongest performance among the tested variants. Removing the gene-aware MoE value embedder reduced performance, consistent with the need to encode expression magnitude in a gene-specific context. Removing spectral subspace projection impaired embedding stability, and removing the symmetric bilinear module produced the largest drops in several benchmarks, indicating that explicit cell-gene coupling is central to the learned biological signal. Variants trained without spatial transcriptomic pretraining also performed worse across multiple settings, supporting the role of spatial data as structural supervision rather than as an optional downstream modality.

### 2.2. OmniCell learns transferable cell representations across species, batches and rare states

A useful cell representation should remain biologically coherent outside the distribution in which it was most directly learned. We therefore benchmarked Omni-Cell against scGPT, scFoundation and Geneformer across human blood and kidney datasets, as well as mouse brain and muscle datasets. OmniCell achieved the highest clustering concordance by both adjusted Rand index and normalised mutual information (Fig. 2a). This advantage was not limited to human data: after homologous gene projection, OmniCell preserved clear cell-type structure in mouse datasets despite being pretrained on human profiles. Representative projections from HumanBlood and MouseMuscle showed compact within-type organisation and sharper separation between related populations than the baseline models (Fig. 2b,c), with the same pattern observed in MouseBrain and HumanKidney (Supplementary Fig. 2a,b). These results are consistent with the role of the soft-rank transformation: by converting raw expression values into within-profile soft ranks, OmniCell preserves relative expression order while reducing sensitivity to sequencing depth, extreme counts and platform-specific magnitude differences, thereby supporting cell embeddings that transfer across tissues and species rather than overfitting to a single atlas.

**Figure 2:**
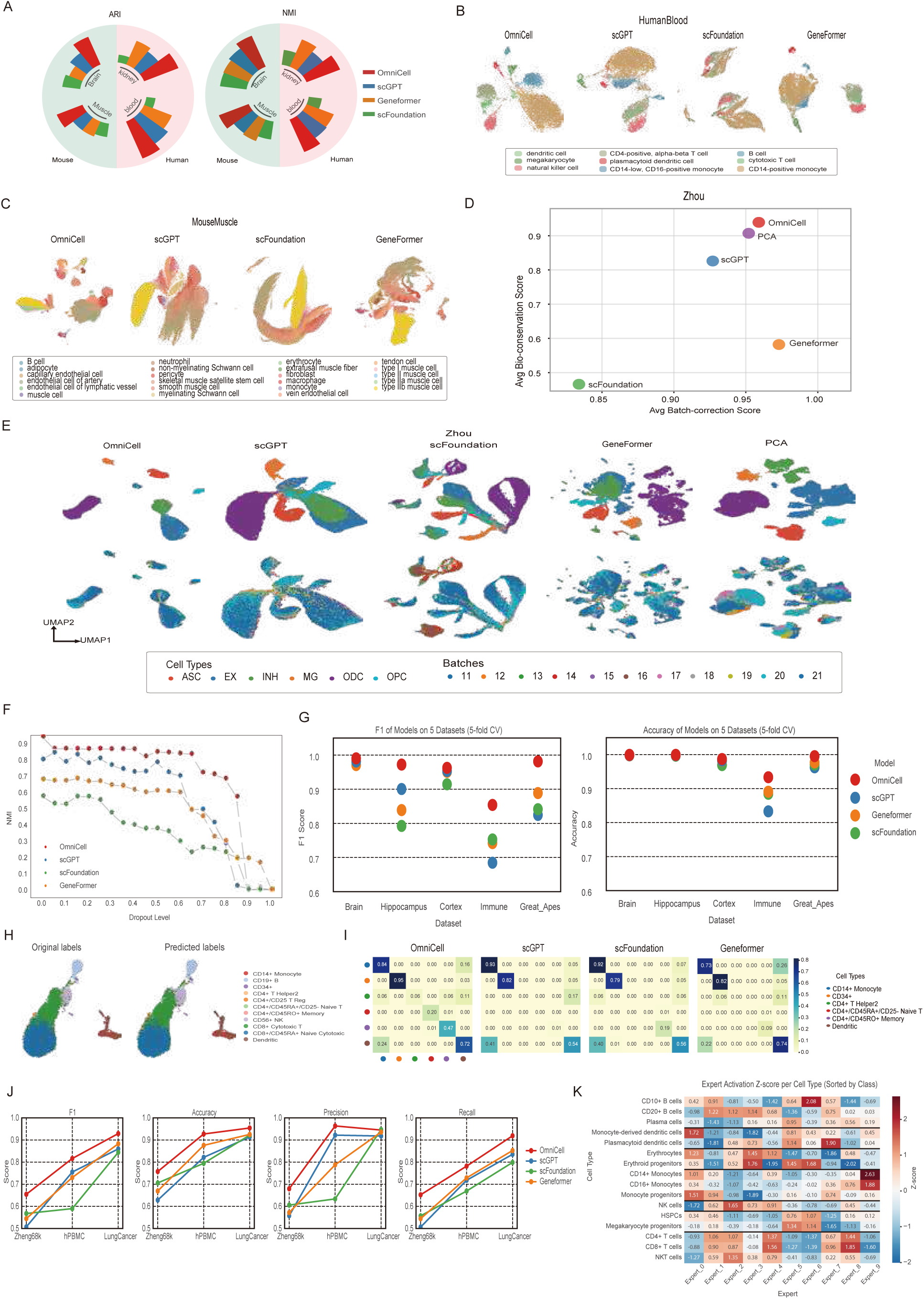
OmniCell learns transferable and robust cell-state representations. (**A**) Cross-dataset clustering performance for OmniCell, scGPT, scFoundation and Geneformer, measured by adjusted Rand index (ARI) and normalised mutual information (NMI) across human blood and kidney datasets and mouse brain and muscle datasets. (**B**,**C**) Representative UMAP projections for HumanBlood and MouseMuscle, showing cell-type organisation produced by each model. (**D**) Batch-integration benchmark on the 11-batch Zhou dataset, comparing average batch-correction score with average biological-conservation score. (**E**) Zhou UMAP embeddings for OmniCell and baseline methods, coloured by annotated cell type (top row) and batch (bottom row). (**F**) Clustering stability under progressive random dropout on the Zhou dataset, quantified by NMI against reference labels. (**G**) Five-fold cell-state annotation performance across Brain, Hippocampus, Cortex, Immune and Great_Apes benchmarks, reported as macro-F1 score and accuracy. (**H**) Zheng68k rare-cell UMAPs comparing original labels with OmniCell-predicted labels. (**I**) Zheng68k rare-class prediction matrices for OmniCell and baseline models, highlighting recovery of low-frequency cell states. (**J**) Overall F1, accuracy, precision and recall across Zheng68k, hPBMC and LungCancer annotation benchmarks. (**K**) Expert-activation z-scores from the gene-aware value embedder across human immune cell types, sorted by lineage group.

The same representation remained stable under technical variation. On the 11-batch Zhou dataset^34^, OmniCell occupied the favourable region of the scIB score space, combining strong batch mixing with preservation of biological conservation (Fig. 2d). Its UMAP embedding mixed cells across batches while retaining the major neuronal and glial populations (Fig. 2e), and a separate hPBMC analysis reproduced this balance across two batches (Supplementary Fig. 2c,d). Progressive random dropout on the Zhou dataset further separated the models: as sparsity increased, OmniCell retained higher clustering stability than scGPT, scFoundation and Gene-former (Fig. 2f). Thus, OmniCell did not improve integration by erasing biological structure. Instead, its embeddings remained informative under both batch variation and increasing sparsity.

Cell-state annotation provided a supervised test of the same embedding space. Across Brain, Hippocampus, Cortex, Immune and Great_Apes benchmarks, OmniCell achieved the strongest macro-F1 score and accuracy under five-fold cross-validation (Fig. 2g). The advantage became most apparent for low-frequency states in Zheng68k. In the main rare-cell view, OmniCell predictions preserved the original UMAP organisation (Fig. 2h), and the rare-class prediction matrices showed clearer diagonal recovery than those of the comparison models (Fig. 2i). The full Zheng68k analysis supported the same pattern across all annotated cell types: the all-class confusion matrix retained stronger diagonal structure and showed broad agreement between original and predicted labels (Supplementary Fig. 2f). Cell-count distributions confirmed that the highlighted Zheng68k classes lay in the low-abundance tail (Supplementary Fig. 2g). CD34+ cells represented only 0.29% of Zheng68k, yet OmniCell identified them with 94% accuracy, 13 percentage points higher than the next-best models. It also recovered CD4+/CD45RA+/CD25− naive T cells with 20% accuracy and CD4+ T helper 2 cells with 6% accuracy, whereas competing models failed to identify these classes (Fig. 2i). Across Zheng68k, hPBMC and LungCancer, OmniCell maintained the strongest overall F1, accuracy, precision and recall profiles (Fig. 2j), with class-level prediction-proportion matrices for hPBMC and LungCancer shown in Supplementary Fig. 2h,i, supporting a consistent gain in both common and rare cell-state annotation.

The gene-aware mixture-of-experts value embedder also provided an interpretable view of the learned cell-state structure. In human immune data, expert activation z-scores followed lineage structure rather than arbitrary model partitions: B cells and plasma cells, myeloid populations, erythroid progenitors, T cells and NKT cells displayed distinct expert usage patterns (Fig. 2k). Expert-preference clustering reproduced this lineage-associated organisation across routing experts (Supplementary Fig. 2e). These results suggest that the routing behaviour reflects biologically coherent cellular programmes inside the model. Together, the cell-level analyses show that OmniCell learns representations that transfer across species and datasets, re-main robust to technical variation, retain rare cellular states and remain interpretable through lineage-aware expert routing.

### 2.3. OmniCell gene embeddings recover cell-type programmes and tissue-aligned modules

OmniCell represents each gene in the context of the cell in which it is observed, so its gene embeddings should carry cell-state information rather than only average expression abundance. We examined this property in Zhou’s human single-nucleus dataset^34^ by extracting cell-level gene embeddings from OmniCell and comparison foundation models. When embeddings for each gene were clustered across cells and compared with annotated cell types, OmniCell achieved the highest adjusted Rand index, indicating stronger preservation of cell-type structure than scGPT, scFoun-dation and Geneformer (Fig. 3a). Gene-specific UMAP projections provided a qualitative view of the same signal: APOE and TREM2 showed expression gradients aligned with cell-type structure in the main analysis, and LHFPL3 showed a similar gene-specific view in the supplementary analysis (Fig. 3b,c; Supplementary Fig. 3c). Attention-derived gene-importance profiles gave an independent readout of cell-type separability: multiclass ROC analysis produced AUC values above 0.9 for all six major cell classes, including astrocytes, excitatory neurons, inhibitory neurons, microglia, oligodendrocytes and oligodendrocyte precursor cells (Fig. 3d).

**Figure 3:**
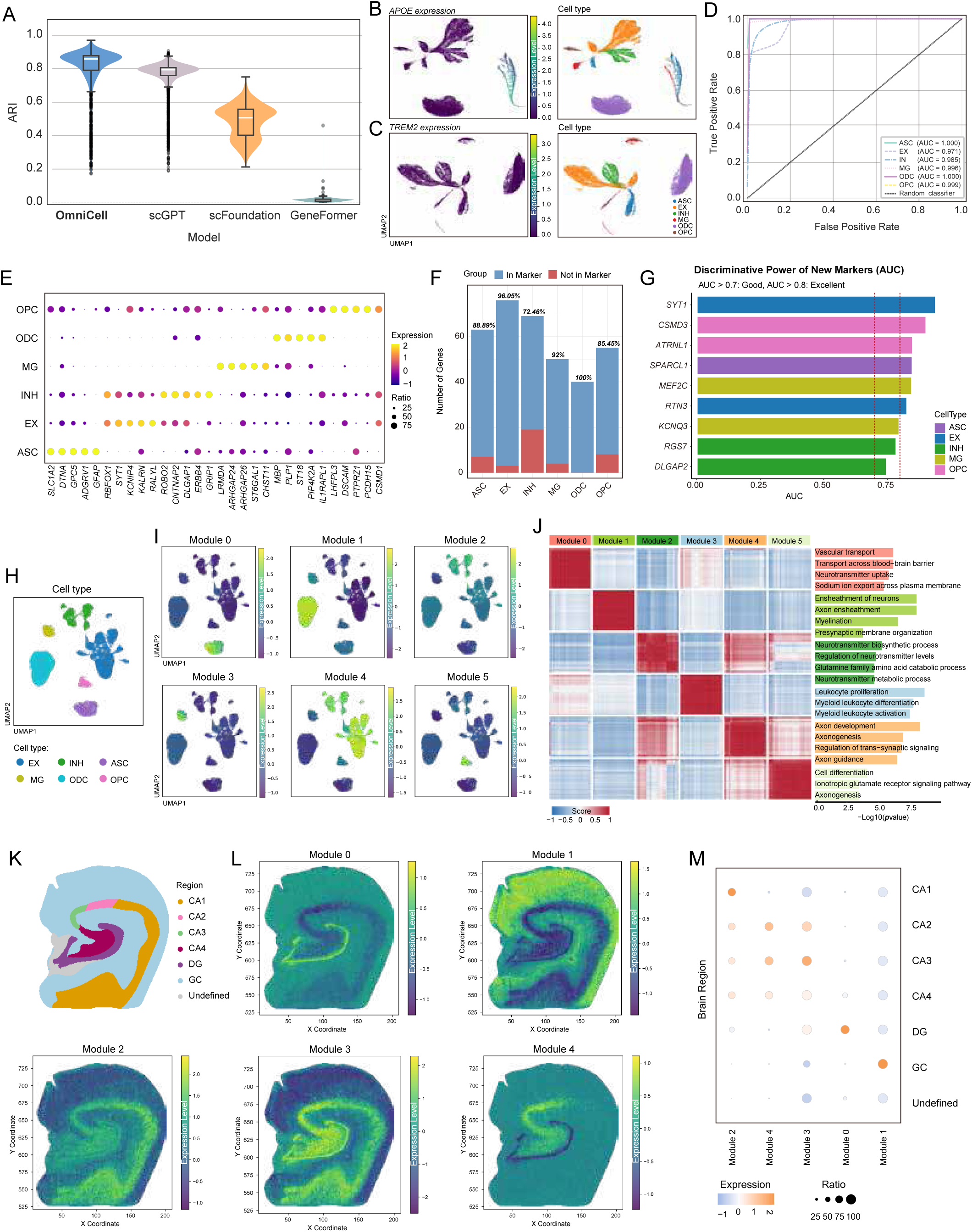
Contextual gene embeddings recover cell-type programmes and tissue-aligned modules. (**A**) Cell-type concordance of gene embeddings in Zhou’s human single-nucleus dataset across OmniCell and baseline models. (**B**,**C**) Cell-level gene-embedding UMAPs for APOE and TREM2; in each panel, the left map is coloured by gene expression level and the right map by annotated cell type. (**D**) Multiclass ROC analysis of attention-derived gene-importance profiles for major neural and glial cell classes. (**E**) Embedding-derived marker genes across astrocytes, excitatory neurons, inhibitory neurons, microglia, oligodendrocytes and oligodendrocyte precursor cells. (**F**) Overlap between embedding-derived markers and expression-ranked markers. (**G**) ROC support for marker candidates not recovered by conventional expression ranking. (**H**) Cell-type reference UMAP for the dataset-level embedding analysis. (**I**) Activity maps for six dataset-level gene co-embedding modules. (**J**) Module score matrix and Gene Ontology enrichment of co-embedding modules. (**K**) Annotated hippocampal regions used for spatial module analysis. (**L**) Spatial activity maps for selected embedding-derived modules in a human hippocampus section. (**M**) Dot plot showing region-level module activity and prevalence across hippocampal annotations.

The same embedding space recovered marker genes with known biological specificity. Ranking genes by cell-type specificity in the OmniCell embedding space and filtering by expression abundance identified canonical markers across major neural and glial populations, including SLC17A7 for excitatory neurons, ERBB4 for inhibitory neurons, GFAP for astrocytes, LRMDA for microglia, PLP1 for oligodendrocytes and PTPRZ1 for oligodendrocyte precursor cells^35–41^ (Fig. 3e; Supplementary Fig. 3b). Most embedding-derived markers overlapped with markers obtained by conventional expression-based ranking, supporting the biological validity of the representation^42^ (Fig. 3f). The non-overlapping candidates were not random residuals: several showed strong discriminative power by ROC analysis, including SYT1 for excitatory neurons and SPARCL1 for astrocytes, each with AUC values above 0.7 (Fig. 3g). Thus, the gene embedding space recapitulated established cell-type programmes while retaining additional marker candidates that remained separable at the representation level.

We then examined whether this structure persisted after moving from cell-level gene embeddings to dataset-level gene relationships. Cell-level gene embeddings were propagated through a cell-similarity graph with a generalised PageRank graph neural network^43^ and then aggregated across cells to obtain dataset-level gene embeddings (Supplementary Fig. 3a). In the Zhou dataset, these dataset-level gene embeddings separated six modules with distinct cell-type activity patterns (Fig. 3h,i). Gene Ontology enrichment supported their functional coherence: the modules were associated with processes such as neurotransmitter transport and synaptic signalling, myelination, immune activation and oligodendrocyte differentiation (Fig. 3j). This module structure indicates that OmniCell gene embeddings do not merely assign individual marker genes, but organise genes into coordinated transcriptional programmes.

Spatial transcriptomic data provided a final test of whether these programmes were aligned with tissue anatomy^29,44^. In human hippocampus samples from Alzheimer’s disease and control tissue^45^, embedding-derived modules showed region-specific spatial activity across cornu ammonis subfields, dentate gyrus and granule cell layer annotations (Fig. 3k–m; Supplementary Fig. 3d,e). Neuron-enriched modules were concentrated in pyramidal cell layers and the dentate gyrus granule cell layer, whereas glial-enriched modules were preferentially localised to the granule cell region. These anatomical patterns link the gene-level representation to spatially organised tissue structure. Together, the Figure 3 analyses show that OmniCell gene embeddings are not a by-product of cell embedding learning, but an interpretable layer that connects cell-type markers, functional gene modules and tissue-aligned programmes.

### 2.4. Spatial representations connect tissue architecture with tumour-margin biology

Spatial transcriptomic interpretation begins with a simple biological requirement: cell identity should remain legible in the native tissue map^2–4^. Across 31 MERFISH mouse brain ageing datasets^46^, OmniCell produced spatial clusters that most closely matched annotated cell types, with NMI and ARI distributions above those of scGPT-spatial and Nicheformer (Fig. 4a). In a representative section, the embedding separated neuronal, glial, vascular and immune populations in UMAP space, and the same labels retained their expected anatomical organisation when projected back onto tissue coordinates (Fig. 4b,c). This slice reached an NMI of 0.8975, compared with 0.8536 for scGPT-spatial and 0.3628 for Nicheformer, and a second MERFISH section showed the same pattern of coherent cell-type separation (Supplementary Fig. 4a). Thus, the spatial signal learned by OmniCell did not blur cell identity; it helped anchor that identity to tissue position.

**Figure 4:**
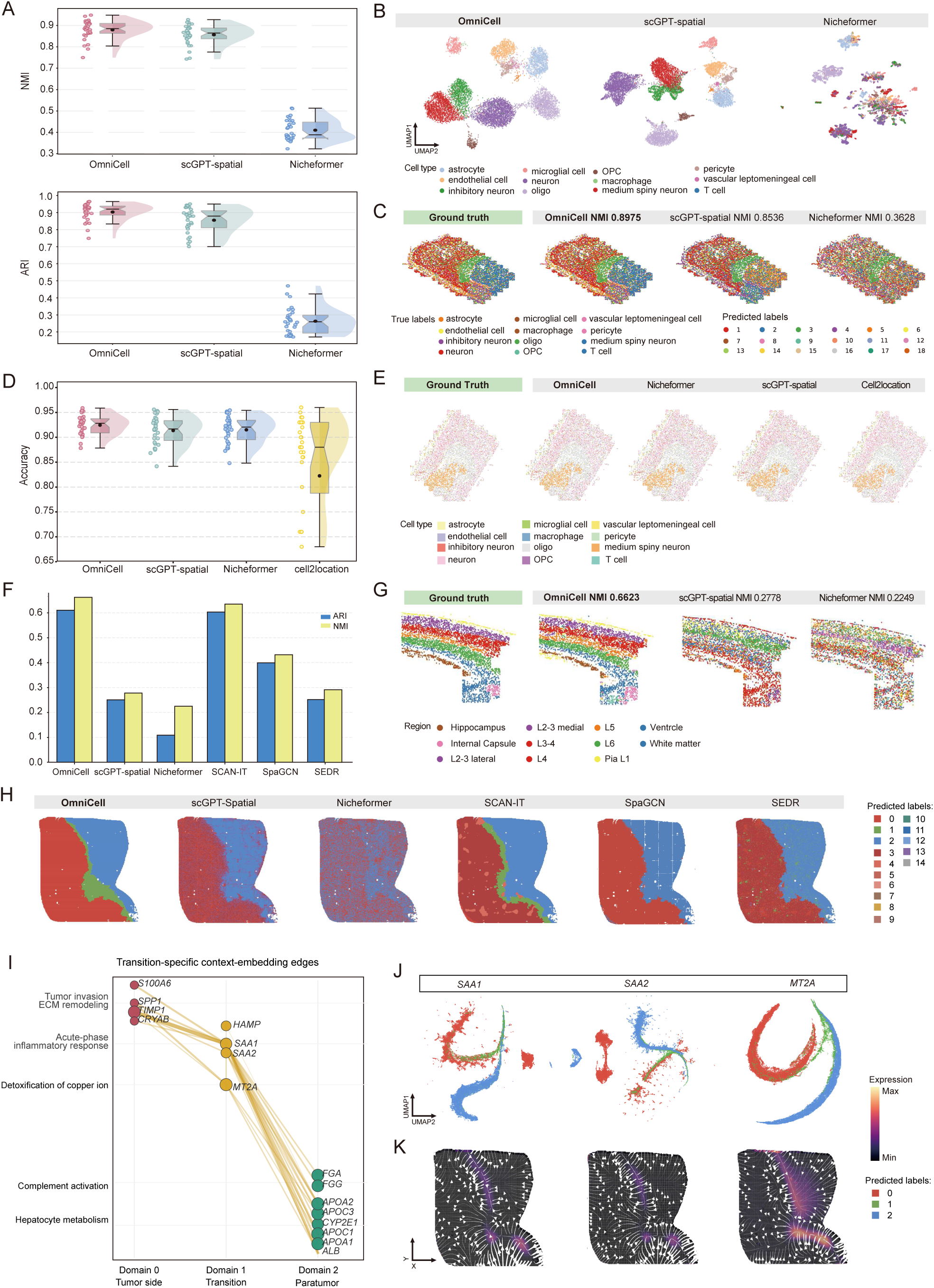
Spatial representations preserve cell identity and resolve tumour-margin transition programmes. (**A**) Spatial clustering performance across 31 MERFISH mouse brain ageing datasets, measured by NMI and ARI for OmniCell, scGPT-spatial and Nicheformer. (**B**,**C**) Representative MERFISH section showing UMAP cell-type separation and tissue-coordinate label maps. (**D**) MERFISH deconvolution accuracy compared with spatial and deconvolution baselines. (**E**) Representative reconstructed cell-type maps relative to annotated reference maps. (**F**,**G**) osmFISH mouse cortex domain reconstruction, including comparison with annotated regions and alternative spatial-domain methods. (**H**) LC5-M liver cancer Stereo-seq spatial segmentation, highlighting a tumour-margin transition zone. (**I**) D1-enriched contextual gene-embedding similarity graph for selected transition-associated genes. Nodes represent genes grouped by their dominant spatial-domain association and functional programme. Edges connect gene pairs whose contextual gene embeddings show higher cosine similarity in the D1 transition context than in D0 or D2. (**J**) Gene-specific spot-embedding UMAPs for SAA1, SAA2 and MT2A, showing the domain structure of each contextual gene embedding. (**K**) Spatial expression maps for SAA1, SAA2 and MT2A projected onto the LC5-M tissue section, highlighting expression along the tumour-parenchyma interface.

Accurate spatial identity should also support a more practical biological task: recovering which cell populations occupy each location^27–28^. In the MERFISH deconvolution benchmark, OmniCell showed the highest and most stable accuracy, whereas cell2location^47^ had a wider performance range and a lower median (Fig. 4d). Reconstructed maps from representative mouse brain slices preserved broad neuronal, glial, vascular and immune distributions that closely followed the annotated reference maps (Fig. 4e; Supplementary Fig. 4c). Confusion matrices for slices 9_0 and 11_2 further showed strong diagonal enrichment for most labelled classes, indicating that the model recovered cell-type composition without collapsing neighbouring but distinct populations (Supplementary Fig. 4d,e).

The same representation extended from cell identity to mesoscale anatomy^28–29^. In an osmFISH mouse cortex section with curated regional annotations^48^, OmniCell reconstructed the laminar and hippocampal structure of the tissue more faithfully than the comparison methods, achieving NMI = 0.6623 and ARI = 0.6108 (Fig. 4f,g). Additional spatial-domain maps for SCAN-IT, SpaGCN and SEDR are shown in Supplementary Fig. 4b. Quantitatively, OmniCell exceeded scGPT-spatial (NMI = 0.2778), Nicheformer (NMI = 0.2249), SpaGCN (NMI = 0.4319), SEDR (NMI = 0.2970) and SCAN-IT^49–51^ (NMI = 0.6354). The strongest visual distinction was the preservation of cortical layers and the separation of white-matter and ventricular regions. These results place the model’s spatial information at a scale between individual cell labels and whole-tissue compartments, where local transcriptomic context helps recover anatomical organisation.

The biological value of this tissue-scale representation became clearest in the LC5-M human liver cancer Stereo-seq dataset, where the relevant structure is not a predefined atlas label but an invasive tumour boundary^1,30^. Tumour margins are biologically active interfaces in liver cancer, where damaged hepatocytes, malignant cells, stromal remodelling and immune suppression can coexist within a narrow invasive zone^30^. OmniCell delineated a continuous transition zone along the tumour-parenchyma interface (Fig. 4h). Among the six tested methods, this border-associated domain was most coherent in the OmniCell segmentation; several baselines either merged the transition with adjacent compartments or broke the section into less interpretable regions.

We next asked whether OmniCell represents gene function as tissue dependent rather than as expression similarity alone. For genes observed across tumour core, transition-zone and paratumour/adjacent non-malignant niches, comparison of contextual gene-embedding cosine similarity across domains revealed clear niche-dependent similarity patterns. The same genes were associated with malignant, proliferative and extracellular-matrix programmes in tumour core, inflammatory and immune stress-response programmes in the transition zone, and hepatocyte metabolic or tissue-homeostatic programmes in paratumour/adjacent non-malignant regions. These shifts match the spatial biology of the liver cancer margin and support the interpretation that OmniCell links gene function to local tissue context.

To test whether this zone carried a coherent molecular programme, we next examined domain-specific contextual gene embeddings rather than raw expression alone. A D1-enriched contextual gene-embedding similarity graph connected transition-associated acute-phase and stress-response genes, including HAMP, SAA1, SAA2 and MT2A, with tumour-side invasion and extracellular-matrix-remodelling genes and paratumour hepatocyte, coagulation and metabolic genes (Fig. 4i). This organisation is consistent with a hepatocyte-injury and inflammatory-interface state. These genes link the transition programme to acute-phase serum amyloid biology, iron homeostasis and metallothionein-mediated metal-ion buffering, processes im-plicated in hepatocellular carcinoma and liver injury^31,52–56^. Gene-specific spot-embedding UMAPs and matched spatial expression maps further localised SAA1, SAA2 and MT2A to the transition interface (Fig. 4j,k). Independent programme enrichment and cell-type composition analyses supported the same interpretation: acute-phase response and copper-detoxification programmes were enriched in D1, and macrophage and T/NK-cell signals increased in the transition zone relative to tumour and paratumour/adjacent non-malignant regions (Supplementary Fig. 4f,l,m). Taken together, these results suggest that OmniCell did not simply recover a geometric boundary. Instead, it resolved a biologically interpretable tumour-margin niche in which hepatocyte injury, acute-phase inflammation, immune-cell accumulation, coagulation/complement activity and metal-ion detoxification converge. We interpret this niche as a testable spatial hypothesis rather than causal proof of the underlying programme.

### 2.5. Virtual perturbations trace developmental timing and cortical-gradient biology

To test whether OmniCell could use tissue context to place counterfactual perturbation responses in biologically appropriate locations, we examined two complementary spatial settings. In the mouse developmental atlas, the question was whether virtual knockouts of well-studied regulators^57–62^ would produce predicted responses at the expected developmental stages and anatomical territories. In the macaque cortex, the question was whether a virtual perturbation could move local transcriptomic states along a published cortical molecular axis^63^. Thus, both analyses tested spatial localisation of predicted perturbation effects rather than experimental validation of gene function.

After fine-tuning on the mouse single-cell spatial transcriptomic atlas^64^, OmniCell separated the three perturbations into distinct developmental trajectories (Fig. 5a–c and Supplementary Fig. 5a). Lhx2 knockout from E12.5 produced a sustained cortical response, with reduced Emx1, Nfix and Pax6 signals concentrated in cortical territories (Fig. 5b–d). Tbr1 knockout from E16.5 shifted the response to a later cortical differentiation programme involving Satb2, Cux2 and Rorb, consistent with disruption of postmitotic cortical identity (Fig. 5b,c and Supplementary Fig. 5b). Foxo1 knockout from P7 instead produced a postnatal striatal phenotype marked by reduced Drd1, Drd2 and Ppp1r1b expression^65^ (Fig. 5b,c,e). Read as a sequence, these predictions trace a developmental story from early cortical regionalisation, through cortical neuronal differentiation, to maturation of striatal projection-neuron states.

**Figure 5:**
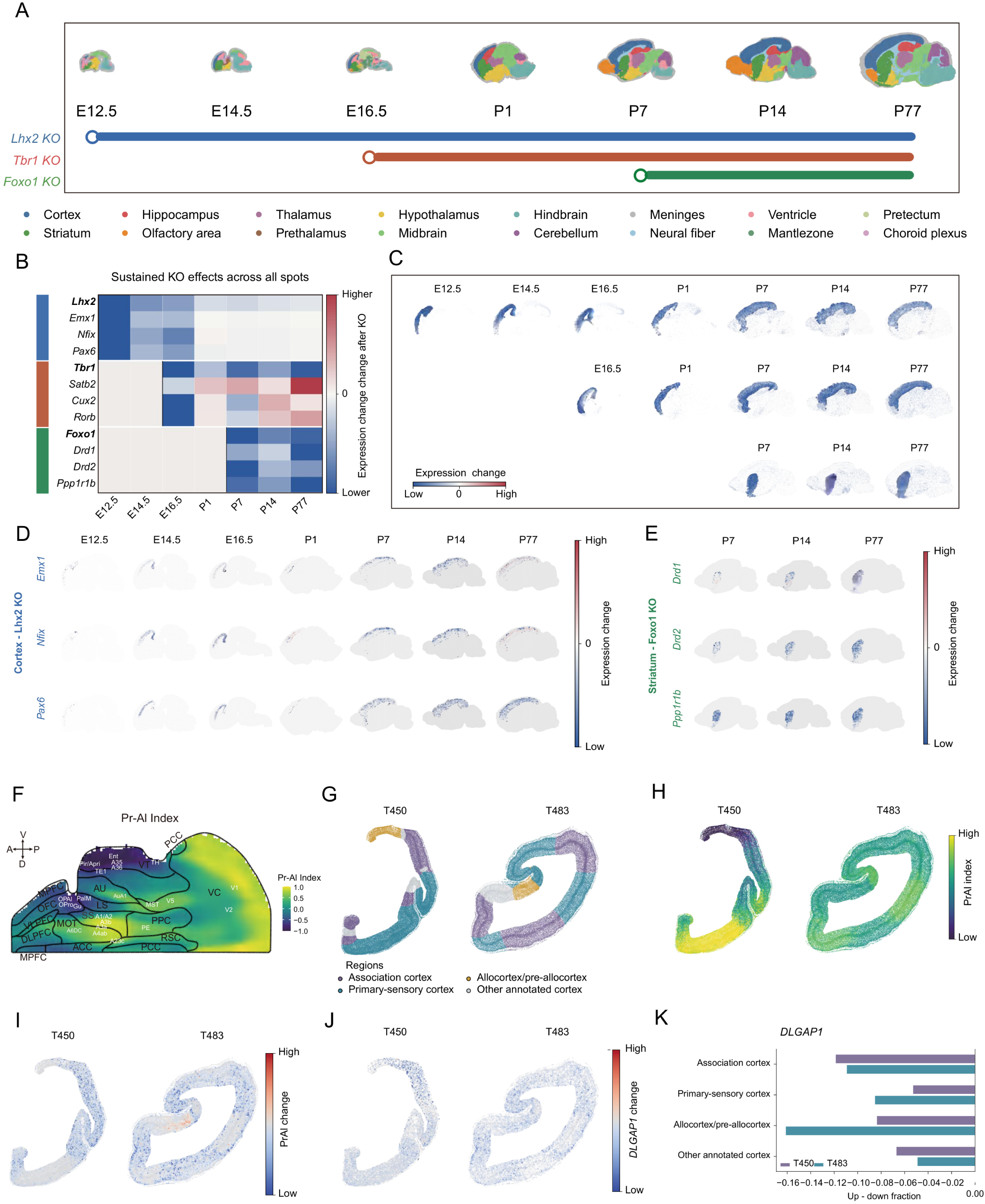
Virtual perturbations predict spatially localised developmental and cortical-gradient effects. (**A**) Mouse developmental spatial atlas and selected virtual knockouts for Lhx2, Tbr1 and Foxo1. (**B**,**C**) Stage- and tissue-resolved knockout effects, separating early cortical, later cortical differentiation and postnatal striatal programmes. (**D**,**E**) Spatial expression-change maps for Lhx2-associated cortical targets and Foxo1-associated striatal targets. (**F**) Published macaque Pr–Al (PRAL) molecular axis used as the cortical-gradient reference; positive values are Pr-like and negative values are Al-like. (**G**) Annotated macaque cortical sections T450 and T483 used for PRAL-based perturbation analysis. (**H**) Baseline OmniCell PRAL index across cortical areas. (**I**) NR3C1 knockout-minus-baseline delta PRAL maps; negative values indicate an Al-shift along the published axis. (**J**) Spatially organised DLGAP1 response after NR3C1 knockout. (**K**) Regional quantification of DLGAP1 response across association, primary-sensory, allocortex/pre-allocortex and other annotated cortical territories.

We then asked whether the same spatial perturbation-prediction framework could move from developmental timing to cortical organisation across species. Omni-Cell was fine-tuned on a macaque spatial transcriptomic atlas in which cortical areas are organised along a primary-sensory-to-allocortex/pre-allocortex molecular axis, referred to here as the Pr–Al (PRAL) Index^63^. In this notation, Al denotes allocortex/pre-allocortex. The PRAL index provides a scalar coordinate between Pr-enriched and Al-enriched transcriptional programmes, so a perturbation can be interpreted as movement along a published cortical organisation axis rather than as a collection of isolated gene changes. We therefore interpreted knockout-minus-baseline decreases in the PRAL index as an Al-shift, indicating movement of the local spatial transcriptomic state toward the Al-like pole; increases would indicate a Pr-shift.

NR3C1 was selected as the primate perturbation target because the reference study identified it as a candidate regulator of the Pr-associated gene programme^63^. If NR3C1 helps maintain Pr-like cortical identity, its virtual knockout should move susceptible regions along the PRAL coordinate. Across two anatomically distant macaque sections, T450 and T483, OmniCell first recovered the expected baseline PRAL organisation, separating primary-sensory cortex, association cortex, allocortex/pre-allocortex and other annotated cortical territories (Fig. 5f–h and Supplementary Fig. 5c). After NR3C1 knockout, delta PRAL maps showed a reproducible Al-shift across sections, most prominently in association cortex, whereas primary-sensory and Al territories showed weaker or more localised displacement (Fig. 5i). The effect therefore appeared as a region-selective displacement along an existing cortical molecular axis, not as a uniform conversion of the cortex.

To connect this gradient-level phenotype to molecular responses, we examined DLGAP1 and DLG1 as concordant NR3C1-associated readouts rather than as independent perturbations. Both genes showed spatially organised expression changes after NR3C1 knockout, and the DLGAP1 response paralleled the region-biased delta PRAL pattern across T450 and T483 (Fig. 5j,k and Supplementary Fig. 5d). Together, the mouse and macaque analyses give the perturbation section a common biological theme: OmniCell did not merely predict that a gene knockout changes expression, but placed the change within a developmental time point, an anatomical territory and a tissue-scale molecular axis. These predictions remain in silico and hypothesis-generating, but they suggest how tissue-contextual foundation models can nominate where a regulatory perturbation may reshape spatial programmes^32–33^.

## 3. Discussion

This study asked whether tissue context can be learned as part of transcriptomic representation, rather than attached after separate single-cell and spatial embeddings have been formed. In this formulation, a gene is not represented only by its average expression pattern, and a cell is not represented only as a dissociated molecular state. Instead, gene activity, cellular state and local tissue neighbourhood jointly define the representation learned during pretraining. OmniCell addresses this question by encoding gene identity, continuous expression values and observed spatial coordinates within one context-aware Transformer, shifting transcriptomic foundation modelling from generic cell-state compression toward tissue-contextual representation learning.

The evidence supports this interpretation across molecular, cellular and tissue scales. At the gene level, OmniCell recovered canonical markers and organised genes into functional modules aligned with cell type and anatomy. At the cell level, it produced embeddings that transferred across species, remained stable under batch variation and simulated dropout, and retained rare cellular states that competing models frequently missed. At the tissue level, it improved spatial clustering, anatomical domain reconstruction and deconvolution. These results indicate that the model is not only compressing high-dimensional count matrices, but learning representations in which molecular state and tissue neighbourhood are mutually informative.

The spatial case studies show why this distinction matters biologically. In the LC5-M Stereo-seq dataset, OmniCell resolved a tumour-margin transition zone marked by macrophage and T/NK-cell accumulation, acute-phase and stress-response programmes, coagulation/complement activity and metallothionein-linked metal-ion detoxification^30–31,52–56^. This result should not be read as proof of a causal mechanism. Rather, it illustrates how a tissue-contextual model can turn representation learning into a testable pathological hypothesis: the invasive tumour boundary may contain a coordinated inflammatory-metabolic niche that is not readily resolved by competing spatial-domain methods. The perturbation analyses extend the same idea to brain development and cortical organisation, where virtual knockouts were mapped onto developmental timing, anatomical territory and the primate PRAL molecular axis. In both settings, the value of the model lies not only in benchmark improvement, but in making spatially structured biological hypotheses visible.

Several limitations define the boundary of these claims. First, representation quality depends on the coverage and quality of the pretraining corpus; extremely sparse spatial datasets, rare cell populations and lowly expressed genes remain challenging. Second, the current training mixture is dominated by dissociated human scRNA-seq and Stereo-seq data, so performance on under-represented tissues, species and spatial technologies may depend on training-set coverage. Third, the perturbation analyses are in silico and hypothesis-generating. Although fine-tuned OmniCell can produce spatial virtual-perturbation predictions for selected regulatory genes, these predictions do not establish experimentally validated causal responses or a general perturbation-response model across all genes, tissues and modalities. This boundary is important because predicting transcriptional responses to chemical or genetic perturbation remains difficult for transcriptomic foundation models^66–68^. Finally, training on tens of millions of profiles requires substantial computational resources, which may limit rapid iteration.

These limitations point to clear next steps. A natural extension is to broaden the pretraining corpus across spatial platforms and species, and to couple transcriptomic tokens with image-derived morphology, perturbation labels or longitudinal measurements. Another is to move from static neighbourhood encoding toward models that explicitly represent developmental progression, immune infiltration or tumour invasion as spatially structured processes. More broadly, the results argue that tissue context should be treated as a first-class axis of foundation modelling. A dissociated cell atlas captures the grammar of cellular identity; adding spatial context begins to capture the tissue syntax in which cellular programmes acquire biological meaning.

## 4. Methods

### 4.1. Pretraining and evaluation datasets

*Pretraining corpus*. Single-cell RNA-sequencing (scRNA-seq) datasets were collected from public CELLxGENE resources, primarily the CZ CELLxGENE Discover Census v.2023-07-15. Datasets associated with this census release were retrieved directly from the publicly accessible Amazon S3 bucket to preserve consistency with the published resource. The final dissociated corpus contained approximately 38 million human single cells across 45 tissues, including brain, blood, lung, liver, digestive and immune systems, skin and kidney. These profiles covered multiple sequencing platforms, developmental stages and disease settings, and accounted for 57.8% of all profiles used for pretraining.

Spatial transcriptomic data were derived primarily from human STOmics Stereo-seq whole-transcriptome datasets generated at BGI Research. The spatial corpus included multiple resolutions, with Bin20 and Cellbin profiles contributing 27.4% and 14.8% of all profiles, respectively. Publicly released Stereo-seq datasets were used when available, and the remaining datasets were drawn from ongoing internal releases. All Stereo-seq profiles were processed with the standard analysis workflow, including spatial binning, UMI quantification and spatial quality assessment. Together, the dissociated and spatial data yielded a pretraining set of approximately 67 million transcriptomic profiles spanning 26 human spatial tissue types.

*Downstream evaluation datasets*. Downstream evaluation was organised to follow the order of the Results. Cell-level clustering and annotation used a diverse set of scRNA-seq datasets, including human kidney datasets^69^, human blood, hPBMC and Zheng68k datasets^5,70–71^, the LungCancer benchmark included in the released benchmark data, mouse brain and muscle datasets, and atlas-level CELLxGENE datasets for Brain, Hippocampus, Cortex, Immune and Great_Apes^72–75^. Batch integration was evaluated on hPBMC and Zhou datasets^34,71,76^. Gene-level analyses used Zhou’s human single-nucleus transcriptomic dataset^34^ and a human hippocampus Stereo-seq dataset containing Alzheimer’s disease and control samples^45^. Spatial analyses used high-resolution spatial transcriptomic datasets, including 31 MERFISH mouse brain ageing samples from the Allen Institute^46^, the LC5-M human liver cancer Stereo-seq dataset^30^, and the osmFISH mouse cortex dataset from the Linnarsson laboratory^48^. Spatial virtual-perturbation analyses used mouse single-cell spatial and macaque cortical spatial transcriptomic atlases^63–64^. Released benchmark data are available at https://modelscope.cn/datasets/PJSucas/OmniCell-test-data.

### 4.2. Unified input representation and sequence construction

*Profile representation*. OmniCell uses one assay-agnostic representation for dissociated and spatial transcriptomic profiles. Gene identities were represented by Ensembl identifiers and mapped to integer indices through a shared vocabulary. For transcriptomic profile *i*, we denote the selected genes by *g_ij_*, their raw expression values by *x_ij_*, and the spatial coordinate of the cell or spot by 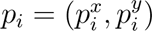. A profile is represented

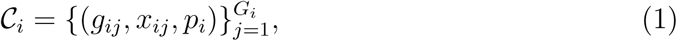

where *G_i_* is the number of selected genes in profile *i*. Spatial transcriptomic profiles retained their measured coordinates. Dissociated scRNA-seq profiles were assigned the placeholder coordinate (0, 0), allowing both modalities to pass through the same context-aware architecture while avoiding any claim that spatial locations were observed for dissociated cells.

*Soft-rank scaling of expression values.* To reduce platform-specific scale effects while preserving within-profile expression order, expression values were transformed by rank-based soft scaling. For each profile *i*, let *u_i_* = {*u_i_*_1_ *< < u_iR_*_i_}be the sorted set of unique selected expression values, with zero included as a reference value when absent from the selected genes. If gene *j* has expression *x_ij_* = *u_irij_*, then its soft rank is

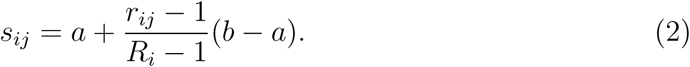

We used *a* = 0 and *b* = 5 throughout. When only one unique value was present, all selected genes were assigned *a*. This transformation maps higher expression to larger soft-rank values, assigns identical expression values the same soft rank, retains relative expression order within each profile, and reduces sensitivity to sequencing depth, extreme counts and platform-specific magnitude differences.

*Sequence construction for dissociated and spatial transcriptomes.* For scRNA-seq, each cell was encoded as an independent gene-token sequence. Non-zero genes were prioritised, followed by high-variance zero-expression genes when padding was needed to maintain sequence length and preserve informative absence patterns. For spatial transcriptomics, each sequence was centred on a focal cell or spot and included its *k* nearest spatial neighbours, selected by Euclidean distance in tissue coordinates; during pretraining, we used *k* = 9, yielding the focal profile plus nine neighbours. The same gene-prioritisation rule was applied to each of the 1 + *k* profiles, selecting *n* genes per profile. Boundary tokens separated neighbouring cellular contexts, enabling the model to distinguish within-cell expression structure from between-cell spatial context while processing the full neighbourhood as one structured input.

### 4.3. OmniCell architecture

*Token embeddings*. Each token embedding combined gene identity with expression magnitude. A shared embedding matrix *E_g_∈*R*^V^* ^×*d*^, where *V* is the vocabulary size and *d* is the model dimension, mapped the integer gene identifier to a dense vector:

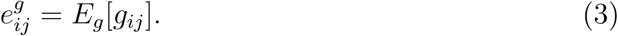

The initial token representation was

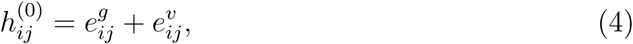

Where 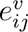 is the expression-value embedding defined below.

*Gene-aware value embedding.* Expression values were embedded with a gene-aware mixture-of-experts (MoE) module. The module used the gene-identity embedding as the routing signal, allowing the same expression magnitude to be encoded differently for different genes. Shared experts captured general abundance trends, whereas routing experts captured gene-conditioned expression behaviour.

For token (*i, j*), routing probabilities over *R* routing experts were computed by a two-layer router with a ReLU nonlinearity:

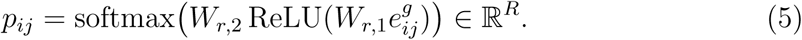

Only the top *K* routing experts were activated:

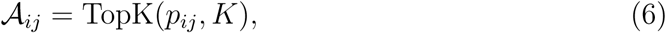

and sparse routing weights were defined without post-selection renormalisation,

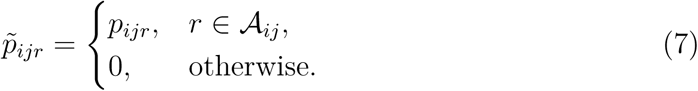

Let 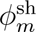 denote the *m*-th shared expert and 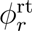 the *r*-th routing expert. The expression-value embedding was

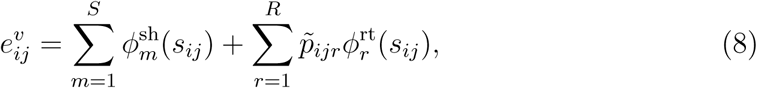

where *S* is the number of shared experts. This design combines global expression-magnitude trends with sparse, gene-conditioned expert specialisation.

*Load balancing for expert use*. To discourage expert collapse, we added a load-balancing regularizer over the routing experts. For a mini-batch containing *T_b_*routed tokens, the empirical selection frequency and mean routing probability for expert *r* were

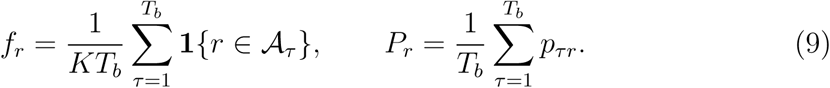

The balancing loss was

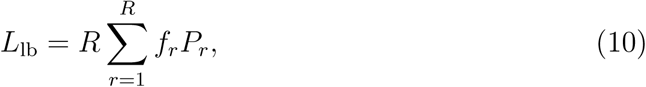

where the factor *R* follows the standard top-*K* load-balancing normalisation. This term encourages both expert-selection frequency and routing confidence to remain broadly distributed across experts.

*Transformer encoder with two-dimensional rotary positional encoding*. The backbone encoder comprised 10 Transformer layers with RMS normalisation, multi-head self-attention, two-dimensional rotary positional encoding (2D-RoPE) and SwiGLU feed-forward networks, for approximately 74 million parameters. Given an input tensor *H*^(0)^ R*^B^*^×*T*^ ^×*d*^, where *B* is the batch size and *T* is the sequence length, each layer applied pre-normalised self-attention followed by a feedforward block.

For a token vector *x* ∈ R*^d^*, RMS normalisation was

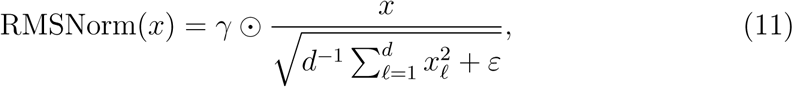

where *γ* is a learned scale vector and *ε* is a numerical-stability constant. In the attention block, query and key vectors were rotated by a block-diagonal rotation matrix *R*(*p*) parameterised by the two-dimensional coordinate *p*. For an attention head,

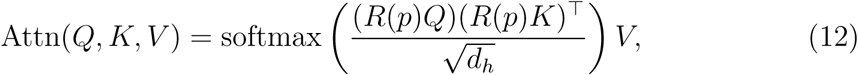

where *d_h_* is the head dimension. Each head was divided into four-dimensional sub-spaces, with paired two-dimensional planes assigned to the *x*- and *y*-coordinate axes. Log-spaced angular frequencies were used for the rotary transformations. This de-sign places tissue coordinates directly inside attention while retaining the sequence representation of gene expression.

The feedforward block used a Swish-gated linear unit,

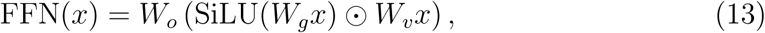

where SiLU(*u*) = *uσ*(*u*), *σ* is the sigmoid function and denotes element-wise multiplication.

### 4.4. Symmetric bilinear output module and pretraining objective

OmniCell was pretrained by soft-rank regression rather than discrete expression-bin prediction. Final-layer token representations were pooled within each semantic cell to produce a cell-level context vector, and a trainable symmetric bilinear trans-formation coupled this context back to token-level features. This objective preserves expression continuity and relative ordering while remaining robust to moderate pre-diction errors.

Let *h_cℓ_* be the final hidden state of the *ℓ*-th interior token in semantic cell *c*. Semantic cells are delimited by boundary tokens; in our implementation, *C* = 1 for scRNA-seq profiles and *C* = 10 for spatial transcriptomic neighbourhoods. The pooled context representation for semantic cell *c* was

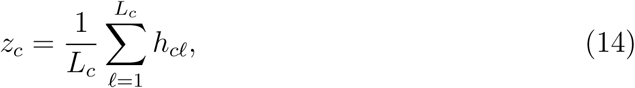

where *L_c_* is the number of interior gene tokens. With a trainable symmetric matrix *M* = *M* ^⊤^, the predicted soft rank for token *ℓ* in semantic cell *c* was

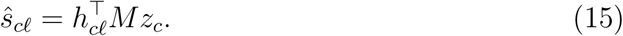

Let Ω denote the set of masked positions and *s_cℓ_* the corresponding target soft ranks derived from observed expression values. The reconstruction loss was

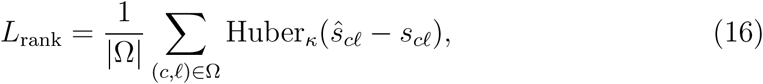

where

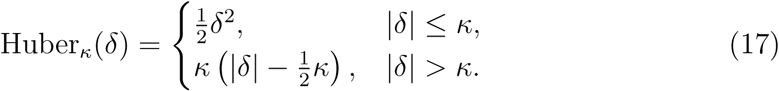

The Huber threshold *κ* was fixed across experiments. The full pretraining objective combined soft-rank reconstruction and MoE load balancing:

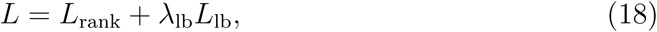

with *λ*_lb_ = 0.05 unless otherwise specified.

### 4.5. Two-stage pretraining

All scRNA-seq and spatial transcriptomic datasets were converted from AnnData (.h5ad) files into LMDB format for high-throughput streaming during large-scale training. Pretraining was performed in two sequential stages. In the first stage, the model was trained on spatial transcriptomic data to learn neighbourhood-dependent co-variation and local tissue continuity. Because spatial profiles are typically sparser and have lower molecular capture efficiency, non-zero expression values in focal cells were masked at a rate of 0.3, non-zero expression values in neighbouring cells were masked at a rate of 0.05, and most zero entries were retained.

In the second stage, the model was further optimised on scRNA-seq data, which provide cleaner single-cell transcriptional signal. Non-zero entries were masked at a rate of 0.3 and zero entries at a rate of 0.1, encouraging the model to infer missing expression from broader transcriptomic context.

Training used a batch size of 3,200 profiles per step, distributed evenly across 32 NVIDIA A100 GPUs with 40 GB memory each. Optimisation used AdamW with a learning rate of 1 × 10^−6^, weight decay of 0.01, momentum coefficients *β*_1_ = 0.9 and *β*_2_ = 0.99, and gradient clipping with maximum norm 1.0. A cosine annealing scheduler with a minimum learning rate of 5 × 10^−7^ was used to support stable convergence.

### 4.6. Inference-time construction of cell and gene embeddings

*Cell embeddings*. Cell embeddings were obtained by aggregating final-layer hidden states with weights that emphasise expressed genes. For cell *i*, the initial aggregated representation was

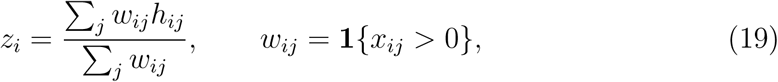

with all selected genes used when no non-zero token was available after filtering. To reduce technical noise, the aggregated vector was optionally projected into a spectral subspace derived from the symmetric bilinear matrix learned during pretraining. If

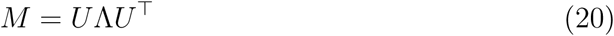

is the eigendecomposition of *M*, the leading basis *U_m_* was selected by cumulative absolute spectral energy. The projected cell embedding was

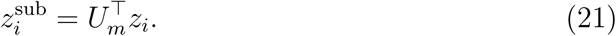

This projection suppresses noise-dominated directions while preserving the dominant biological structure captured by the bilinear output module.

*Cell-level and dataset-level gene embeddings*. Cell-level gene embeddings were taken from the final hidden state of each gene token, followed by the same optional spectral projection. These contextualised vectors represent how each gene behaves in a specific cell state. Dataset-level gene embeddings were obtained by propagating cell-level gene vectors over a cell-similarity graph (Supplementary Fig. 3a). The graph was constructed from OmniCell cell embeddings by connecting each cell to its nearest neighbours and weighting edges with a decaying kernel. For each gene, its cell-level embeddings were propagated with a generalised PageRank graph neural network^43^ and then averaged across cells. This procedure reinforces co-expression patterns and context-dependent gene relationships across the full dataset.

### 4.7. Cell-level analyses

*Single-cell clustering across species*. Cell clustering was evaluated for OmniCell, scGPT, Geneformer and scFoundation on human and mouse scRNA-seq datasets. For mouse datasets, gene identifiers were first projected onto the human gene vocabulary through homologous gene mapping before model inference. We retained one-to-one mouse–human homologous gene pairs and discarded mouse genes without a unique human counterpart or without a corresponding entry in the model vocabulary, so each retained mouse expression feature was represented by the matched human gene identifier used during pretraining. The top 2,000 highly variable genes were then selected from the mapped gene space for all scRNA-seq datasets. Embeddings were clustered with the Leiden algorithm across resolutions from 0.1 to 1.0, and the resolution with the highest agreement with reference labels was selected. Performance was quantified with ARI and normalised mutual information (NMI).

*Batch correction and dropout robustness*. Batch-correction performance was evaluated with the scIB benchmarking framework^77^. PCA, scGPT, Geneformer and sc-Foundation were used as baselines. hPBMC^71^ and Zhou^34^ datasets were evaluated with the top 2,000 highly variable genes. The scIB biological conservation score measured preservation of annotated cellular structure, whereas the batch-correction score measured integration of technical batches by silhouette and graph-connectivity metrics.

Noise robustness was tested on the Zhou dataset by introducing random dropout noise from 0.0 to 1.0 in increments of 0.05. For each noise level, embeddings were regenerated and clustered with Leiden clustering at resolution 0.1. NMI against reference annotations was used as the primary stability metric. All experiments used fixed random seeds.

*Cell-type annotation and rare-state recovery*. For supervised annotation, the top 2,000 most variable genes were selected from each dataset and model-derived cell embeddings were used as classifier inputs. OmniCell was compared with scGPT, Geneformer and scFoundation. A three-layer multilayer perceptron was trained with ReLU activation, layer normalisation and AdamW optimisation for 20 epochs at learning rate 1 × 10^−4^. Evaluation used five-fold stratified cross-validation for the Brain, Hippocampus, Cortex, Immune and Great_Apes datasets^72–75^, reciprocal cross-batch validation for hPBMC^71^, and an 80:20 stratified train-test split for large sparse datasets such as Zheng68k^5^ and the LungCancer benchmark. Accuracy and macro-averaged F1 score were reported as mean and variance across folds or batches. Predicted labels and confusion matrices were used to inspect recovery of low-frequency cell states in Zheng68k.

*Expert-routing analysis*. To characterise the cell-type selectivity of MoE experts, we first averaged expert routing weights across cells within each annotated cell type, generating a cell type-by-expert matrix. For each expert, routing weights were then Z-score normalised across cell types to measure relative expert activation. The resulting Z-score matrix was visualised as a heatmap after sorting cell types by predefined biological groups, with group boundaries indicated by horizontal lines.

To further quantify expert preference, cell types with Z-score > 1.0 were identified for each expert. These high-activation cell types were mapped to broader biological groups, and the group with the largest proportion was assigned as the preferred group of that expert. The preference score was defined as the proportion of high-activation cell types belonging to this dominant group. Experts were then grouped by their preferred cell-type category and visualised using a colour-coded bar plot.

### 4.8. Gene-level analyses

*Cell-type specificity of gene embeddings*. To assess whether cell-level gene embeddings preserved cell identity, final-layer gene representations were extracted from OmniCell and from the comparison foundation models. Using the Zhou dataset and 2,000 highly variable genes, embeddings for each gene were collected across cells and clustered in the corresponding embedding subspace. The resulting gene-specific clusters were compared with known cell-type annotations using Adjusted Rand Index (ARI). Gene-cell importance scores were also derived by pooling gene-gene attention weights across Transformer layers. For each cell type, a prototype importance pro-file was computed by averaging importance vectors across cells of that type. Cosine similarity between each cell’s importance vector and the cell-type prototypes yielded scores for multiclass receiver operating characteristic analysis and area-under-the-curve calculation.

*Marker-gene discovery from embedding space*. Cell-type-specific genes were identified by comparing each gene’s cell-level embedding distribution in a target cell type against its distribution in all other cells. Distinctiveness was quantified as the mean one-dimensional Wasserstein distance across embedding dimensions. Genes were ranked by this embedding-space discrepancy and then filtered by transcriptional abundance, retaining genes with higher mean expression in the target cell type than in other cells. The resulting markers were compared with markers obtained from Seu-rat FindAllMarkers^71^. Additional embedding-derived markers that did not overlap with conventional marker lists were evaluated by ROC analysis and AUC.

*Dataset-level module discovery*. To test whether dataset-level gene embeddings captured coordinated transcriptional programmes, gene-gene similarity networks were constructed with cosine similarity or a Gaussian-transformed Euclidean metric. Soft thresholding was applied to enrich high-confidence relationships, and modules were identified by Louvain community detection. Biological relevance was evaluated in the Zhou scRNA-seq dataset^34^ and in Alzheimer’s disease and control hippocampus Stereo-seq data^45^ using cell-type marker genes and region-specific marker genes as priors. Module expression scores were calculated to quantify alignment between embedding-derived modules and transcriptomic co-expression structure.

### 4.9. Spatial analyses

*Spatial cell clustering*. Spatial clustering was evaluated with OmniCell, scGPT-spatial and Nicheformer on the MERFISH mouse brain ageing datasets^46^, for which the full gene set was retained. Spatial-mode embeddings were clustered with the Leiden algorithm across resolutions from 0.1 to 1.0, and the resolution with the highest agreement with reference labels was selected. Performance was quantified with ARI and NMI, and representative embeddings were projected back onto tissue coordinates to inspect anatomical coherence.

*Spatial transcriptomic deconvolution*. Spatial cell-type deconvolution used a Tangram-like alignment strategy^78^. Single-cell reference profiles and spatial transcriptomic profiles were aligned by shared genes. Single-cell embeddings defined the reference space, and spatial embeddings were projected into that space by optimisation to estimate the proportional contribution of each cell type to each spatial location. The dominant cell type at each spot was assigned by the largest estimated contribution. OmniCell was compared with scGPT-spatial, Nicheformer and cell2location^47^, and performance was evaluated with accuracy and macro-averaged F1 score.

*Spatial domain identification and biological characterisation*. Spatial domain reconstruction was evaluated on a mouse cortex dataset with curated anatomical annotations and on the LC5-M human liver cancer Stereo-seq dataset. For mouse cortex, all 33 genes were used during inference. Spatial-mode embeddings were generated by integrating a focal profile with its neighbours, followed by local averaging, construction of a k-nearest-neighbour graph and Leiden clustering. The clustering resolution was optimised from 0.1 to 1.0 in increments of 0.1 by maximising agreement with the reference annotations.

For LC5-M, OmniCell used the top 500 highly variable genes, consistent with the pretraining configuration, whereas baseline methods used their standard top 2,000 genes. Spatial-mode embeddings were clustered with a k-nearest-neighbour graph and Leiden resolution 0.1. OmniCell was compared with scGPT-spatial, Nicheformer, SpaGCN, SEDR and SCAN-IT^49–51^. Domain-specific programme enrichment, D1-enriched contextual gene-embedding similarity graph analysis, gene-specific spot-embedding visualisation, matched spatial expression mapping and cell-type composition analysis were used to characterise spatial domains identified in the tumour, transition-zone and paratumour/adjacent non-malignant compartments.

### 4.10. Spatial virtual-perturbation analyses

Spatial virtual-perturbation analyses were designed to test whether a fine-tuned OmniCell model could localise the predicted effect of selected regulatory perturbations within an anatomical map^79^. For mouse development, OmniCell was fine-tuned on the mouse single-cell spatial transcriptomic atlas^64^ and used to compare baseline predictions with virtual knockouts of Lhx2, Tbr1 and Foxo1. These genes were selected because prior studies place them in early cortical patterning, postmitotic cortical differentiation and striatal projection-neuron development, respectively^57–62^. Perturbation effects were summarised as knockout-minus-baseline changes in stage-and region-relevant marker genes, including Emx1, Nfix and Pax6 for the Lhx2 analysis; Satb2, Cux2 and Rorb for the Tbr1 analysis; and Drd1, Drd2 and Ppp1r1b for the Foxo1 analysis^65^.

For primate cortical analysis, OmniCell was fine-tuned on a macaque spatial transcriptomic atlas with published primary-sensory-to-allocortex/pre-allocortex organisation^63^. The reference study defined an opposing Pr–Al (PRAL) molecular axis by comparing each cortical region or segment with primary sensory (Pr) and allocortex/periallocortex (Al) references. For macaque data, the Pr reference comprised areas 3a/b (S1), V1 and AuA1, whereas the Al reference comprised piriform cortex and entorhinal subdivisions ER, EO, EI, ELr, ELc, ECL and EC^63^. Pearson correlations to the Pr and Al references were summarised as Cor_Pr_ and Cor_Al_, independently normalised by setting the maximum, mean and minimum to 1, 0.5 and 0, respectively, and combined as

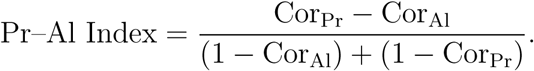

Positive values therefore indicate Pr-like cortical composition, whereas negative values indicate Al-like composition. We applied this published PRAL index as a scalar summary of the cortical molecular axis rather than deriving a new gradient. NR3C1 was selected as the perturbation target because the reference study identified it as a candidate regulator of the Pr-associated gene programme. Baseline and NR3C1-knockout predictions were compared across the T450 and T483 sections, and perturbation effects were reported as knockout-minus-baseline delta PRAL maps and as spatial expression changes in DLGAP1 and DLG1. Negative delta PRAL values were interpreted as an Al-shift along the published axis. These analyses were interpreted as computational hypotheses rather than experimental validation of causal gene function.

### 4.11. Baseline implementations

Foundation-model baselines were run with official pretrained weights and inference pipelines. scGPT embeddings were generated from the scGPT_human pretrained model. Geneformer embeddings were generated with Geneformer-V2-316M from the official Hugging Face repository, using penultimate-layer outputs and mean pooling of gene-level embeddings into cell-level representations. scFoundation used the published models.ckpt checkpoint and the authors’ standard inference procedure. scGPT-spatial used the scGPT_spatial_v1 checkpoint pretrained on spatial transcriptomics. Nicheformer used released parameters trained on SpatialCorpus-110M. Specialised spatial-domain methods, including SpaGCN v1.2.7, SEDR and SCAN-IT^49–51^, were run with official implementations and recommended settings unless dataset-specific preprocessing required otherwise.

## Data availability

No new raw sequencing data were generated in this study. Public datasets analysed in this study are available from the original sources cited in the Methods and References.

## Code availability

OmniCell is distributed as an open-source repository at https://github.com/BGIResearch/omnicell.

## Acknowledgements

We thank the Stomics Cloud platform (https://cloud.stomics.tech/) for providing GPU computational resources. We also thank our research group colleagues for insightful discussions and contributions.

## Author contributions

X.X. conceptualised the study. P.J.S. was responsible for framework design and tool implementation. P.J.S., Q.P., D.Y.T., H.Y.Z., L.B.L. and T.W.Y. performed data analysis and model evaluation. L.B.L. and T.W.Y. were responsible for building the baseline model. T.F., L.A.D., C.L., H.Z., Y.J., L.S.K., D.Z.Q. and F.S.S. provided key suggestions. P.J.S., Q.P. and H.Y.Z. wrote the manuscript. Z.Y., L.Y.X. and L.S.S. supervised the study.

## Competing interests

The authors declare no competing interests.

---

**Supplementary Figure 2.**
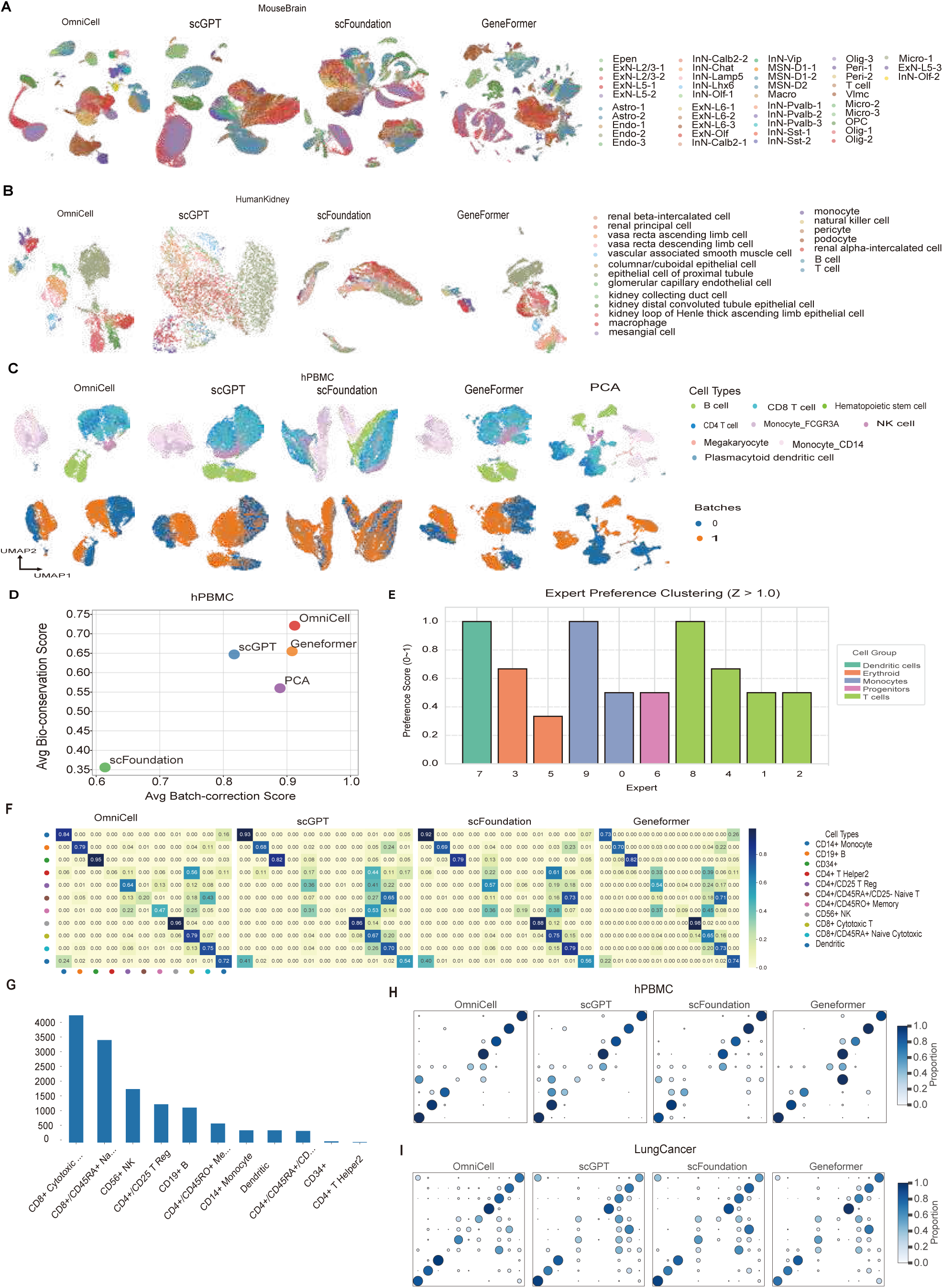
Additional cell-representation benchmarks. (**A**) MouseBrain UMAP projections for OmniCell, scGPT, scFoundation and Geneformer, coloured by annotated cell type. (**B**) HumanKidney UMAP projections for the same models, coloured by annotated cell type. (**C**) hPBMC UMAP embeddings for OmniCell, scGPT, scFoundation, Geneformer and PCA, coloured by annotated cell type (top row) and batch (bottom row). (**D**) hPBMC batch-integration benchmark, showing average batch-correction and biological-conservation scores. (**E**) Expert-preference scores for high-activation routing experts (standardised activation *Z >* 1.0); bar height denotes the preference score and colour denotes the broad cell group associated with each expert. (**F**) Zheng68k all-class row-normalised confusion matrices across models. (**G**) Cell-count distribution of Zheng68k annotated classes used to identify low-frequency states. (**H**) hPBMC class-level prediction-proportion dot matrices across models. (**I**) LungCancer class-level prediction-proportion dot matrices across models.

**Supplementary Figure 3.**
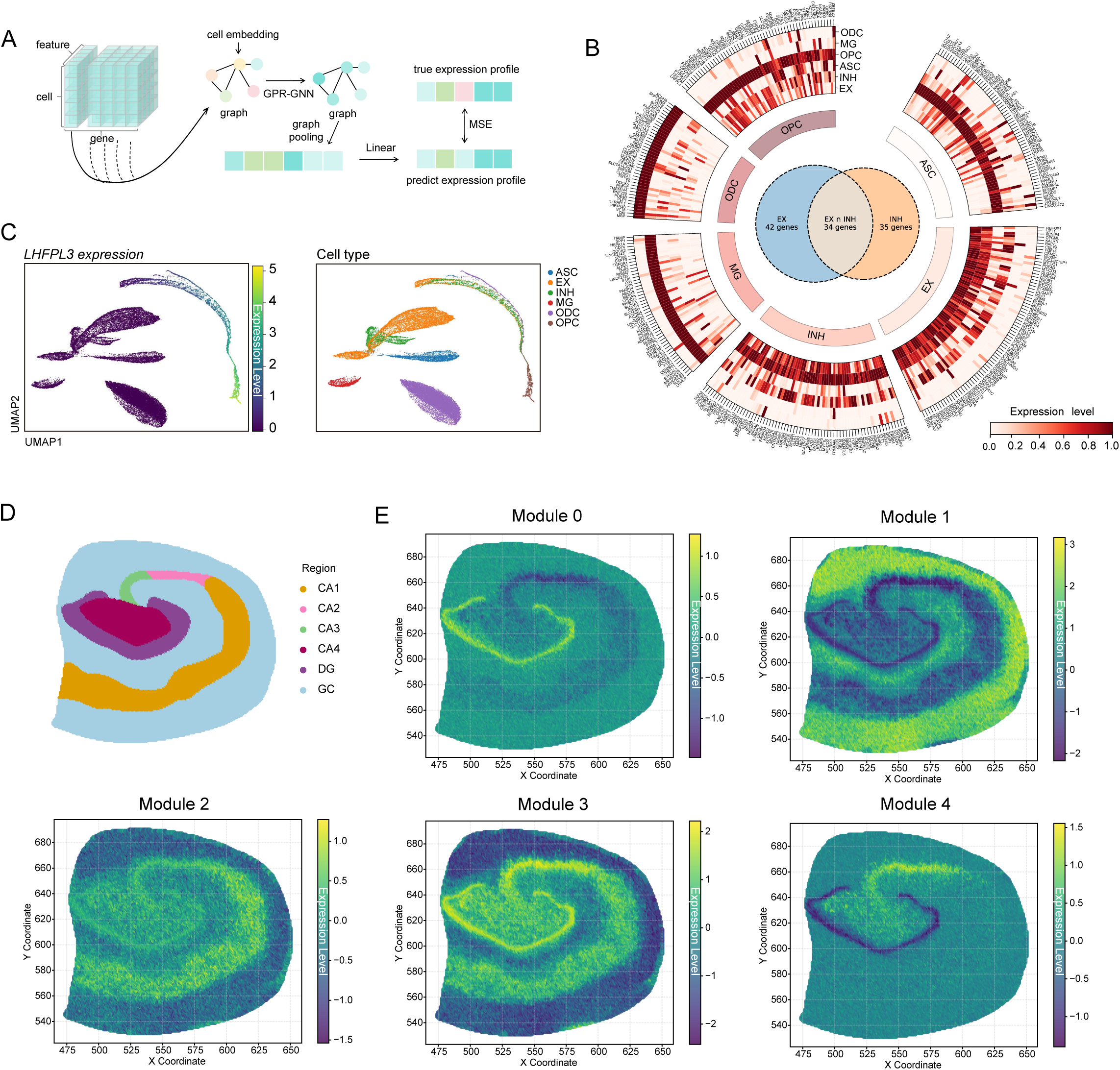
Additional gene-embedding and spatial-module analyses. (**A**) Graph-propagation workflow used to derive dataset-level gene embeddings from cell-level gene features and cell-similarity structure. (**B**) Embedding-derived marker-gene view across astrocytes, excitatory neurons, inhibitory neurons, microglia, oligodendrocytes and oligodendrocyte precursor cells. (**C**) Cell-level gene-embedding UMAP for LHFPL3; the left map is coloured by LHFPL3 expression level and the right map by annotated cell type. (**D**) Annotated hippocampal regions used for spatial module analysis. (**E**) Spatial activity maps for embedding-derived modules across the hippocampal section.

**Supplementary Figure 4.**
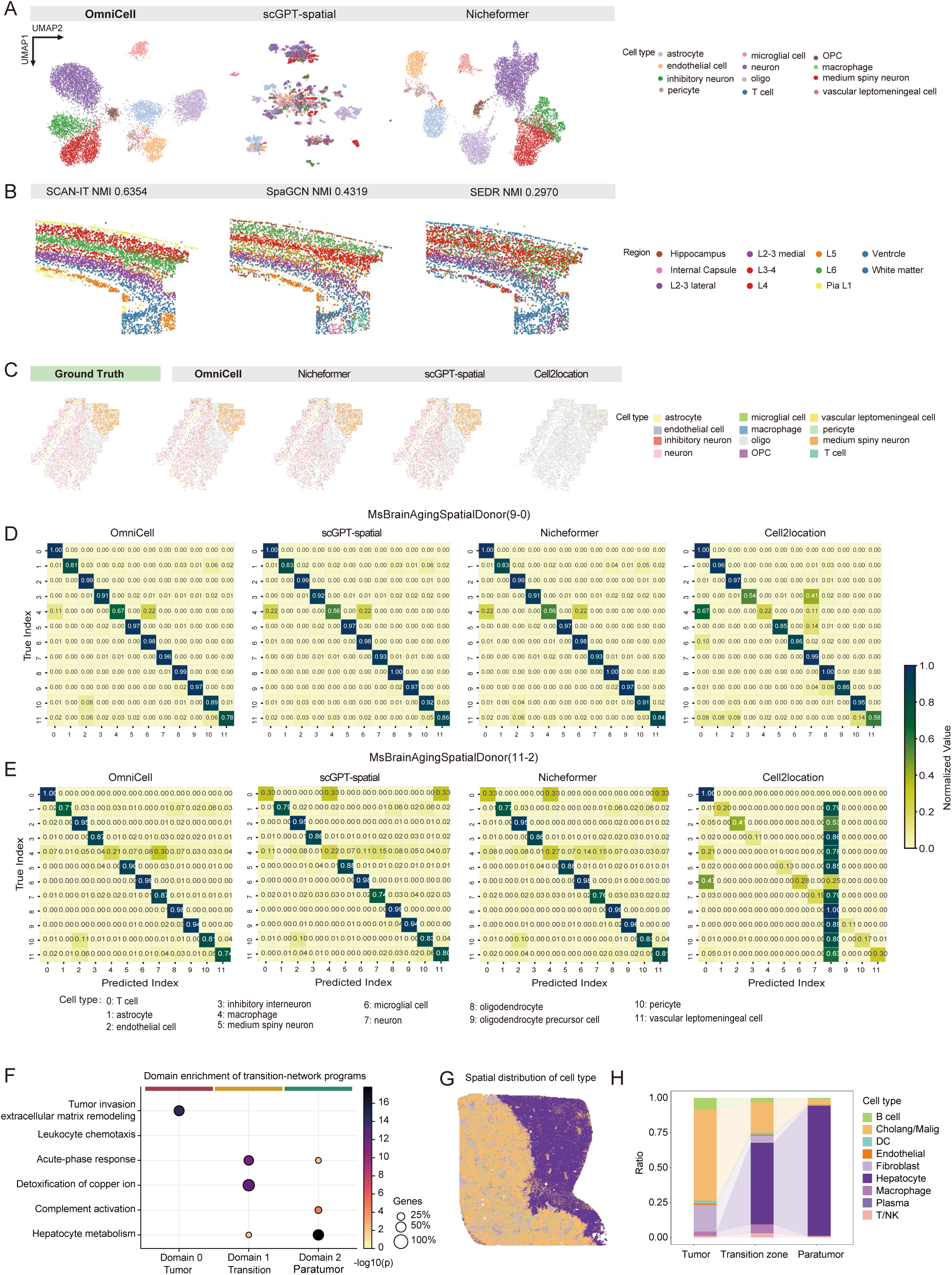
Additional spatial clustering, domain-reconstruction and LC5-M transition-zone analyses. (**A**) UMAP embeddings for a second MERFISH mouse brain section across OmniCell, scGPT-spatial and Nicheformer, coloured by annotated cell type. (**B**) osmFISH domain-reconstruction maps for SCAN-IT, SpaGCN and SEDR, with NMI values shown above each map. (**C**) Representative MERFISH deconvolution maps showing the ground truth and reconstructions from OmniCell, Nicheformer, scGPT-spatial and Cell2location. (**D**) Normalised confusion matrices for MERFISH donor 9_0 across the evaluated models. (**E**) Normalised confusion matrices for MERFISH donor 11_2 across the evaluated models. (**F**) Domain enrichment of transition-network functional programmes in LC5-M. Dot colour indicates enrichment significance, shown as − log_10_ (*P*), and dot size indicates the fraction of genes from each programme represented in the domain. (**G**) Spatial distribution of major cell types in the LC5-M tissue section. (**H**) Cell-type composition across tumour, transition-zone and paratumour/adjacent non-malignant regions.

**Supplementary Figure 5.**
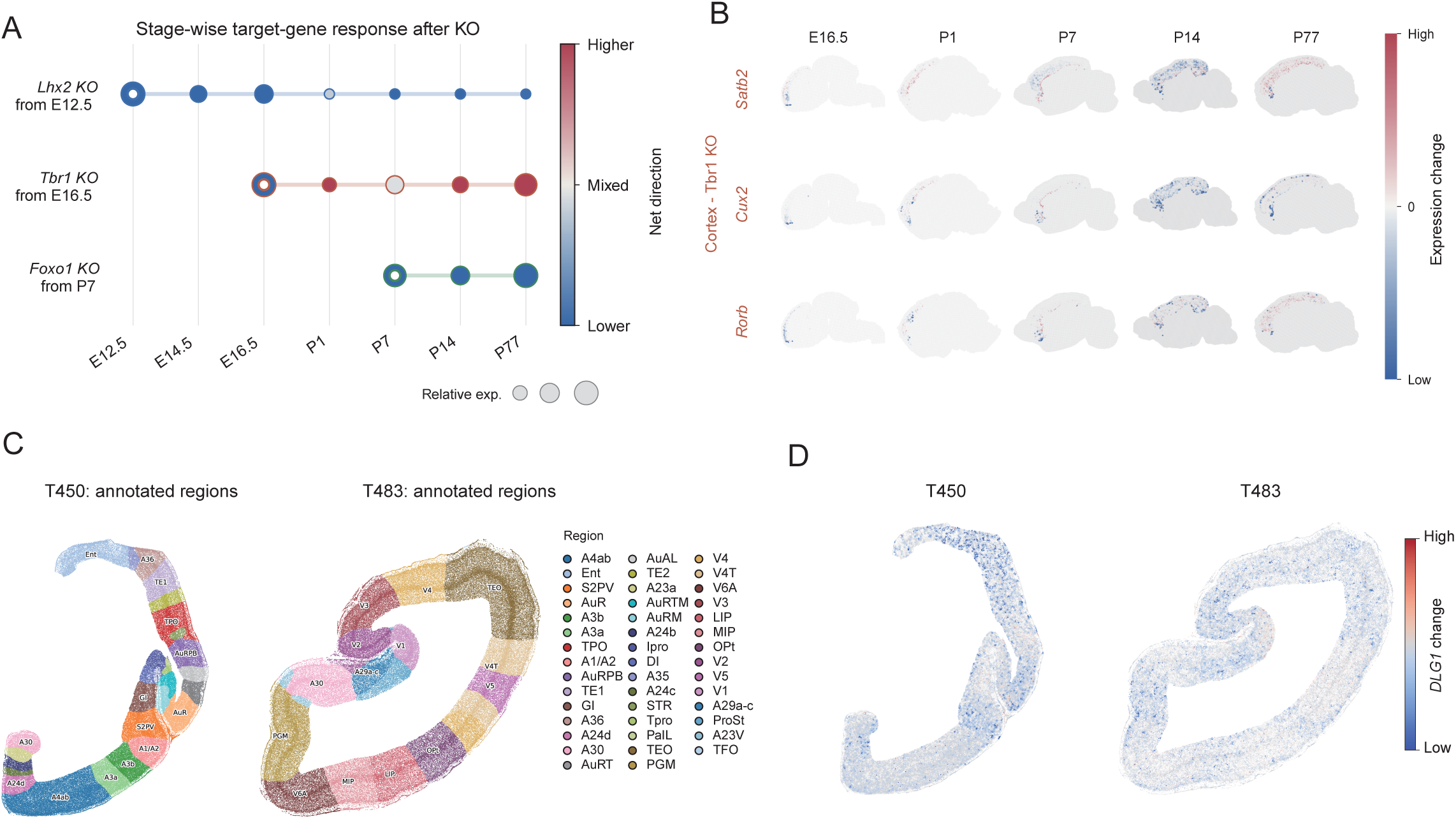
Additional spatial virtual-perturbation analyses. (**A**) Stage-wise target-gene responses after Lhx2, Tbr1 and Foxo1 virtual knockout. (**B**) Spatial expression-change maps for Tbr1-associated cortical differentiation genes across developmental stages. (**C**) Annotated macaque cortical regions in the T450 and T483 sections. (**D**) DLG1 expression-change maps after NR3C1 knockout in T450 and T483.

## Notes

### Competing Interest Statement

The authors have declared no competing interest.

